# Collective phases and long-term dynamics in a fish school model with burst-and-coast swimming

**DOI:** 10.1101/2024.05.26.595998

**Authors:** Weijia Wang, Ramón Escobedo, Stéphane Sanchez, Zhangang Han, Clément Sire, Guy Theraulaz

## Abstract

Intermittent and asynchronous burst-and-coast swimming is widely adopted by various species of fish as an energy-efficient locomotion mode. This swimming mode significantly influences the way in which fish integrate information and make decisions in a social context. Here, we introduce a simplified fish school model in which individuals have an asynchronous burst-and-coast swimming mode and selectively interact only with one or two neighbors that exert the largest influence on their behavior and over a limited spatial range. The interactions consist for a fish to be attracted and aligned with these neighbors. We show that, by adjusting the interactions between individuals above a sufficiently high level, depending on the relative strength of attraction and alignment, the model is able to produce a cohesive fish school that replicates the main collective phases observed in nature: schooling, milling and swarming when each individual interacts with only one neighbor, and schooling and swarming when each individual interacts with two neighbors. Moreover, the model shows that these patterns can be maintained over long simulations. However, with the exception of swarming, these patterns do not persist indefinitely in time and fish lose cohesion and progressively disperse. We further identify the mechanisms leading to the dispersion.

---

Many species of fish living in groups adopt an intermittent “burst-an-coast” swimming mode [1, 2]. This way of swimming is characterized by an active swimming “burst” phase followed by a passive gliding phase and plays many roles in the life of fish, such as energy saving [3–6], decrease of detectability [7] and the stabilization of the sensory field [2].

When moving in groups, fish coordinate their swimming through their social interactions: they are both attracted to their neighbors and also align with them [8– 10]. The form of these social interactions varies between species and depends on the distance between fish as well as on the orientation and relative position of their neighbors [11–13]. For instance, the range of attraction and alignment interactions has been shown to be much larger in *H. rhodostomus* than in *D. rerio* [14]. Furthermore, several studies have shown that each fish does not interact with all its neighbors but only with a small subset of them [15–17], and, in particular, with those that exert the strongest influence on its movement [18].

These most influential individuals are those that at each moment trigger a larger response than other neighbors. This selection reduces the amount of information that needs to be processed in the fish’ brain and avoids cognitive overload. Finally, individuals are able to modulate the intensity with which they interact with their neighbors according to their physiological state or the characteristics of the environment [19].

Understanding how the combination of these social interactions between fish and their modulation by physiological or environmental parameters determine the types and properties of collective movements at the level of a school is a fundamental question in the research field of collective animal behavior [20–23]. Most existing fish school models consider that individuals have a continuous swimming mode. Very few models have studied the impact of asynchronous and intermittent movement on swimming coordination, and even fewer have explored its impact on the resulting collective movement phases [24– 27].

In this article, we introduce a general model to study the properties of collective movements of fish schools in which individuals have an asynchronous and intermittent swimming mode. This model is a simplified version of a data-based model initially developed to account for collective swimming behavior in *Hemigrammus rhodostomus* [13, 18, 27]. The model integrates both the intermittent swimming mode of fish, the asynchronous individual decisions at the onset of the bursting phase, as well as the filtering of social information which results from the fact that each fish only interacts with at most its two most influential neighbors.

We first describe the model and we analyze the different collective phases produced by groups of fish swimming in an unbounded environment for different combinations of the intensity of the attraction and the alignment interactions. We then analyze the long-term behavior, the stability of each collective phase, and the mechanisms by which this stability is eventually lost in the long term.

We show that this simplified burst-and-coast model in which fish interact asynchronously with only one or two neighbors and within a limited perception range is able to reproduce the main features of the original data-based model [27] in particular a cohesive fish school and the typical collective phases of swarming, schooling, and milling. Moreover, the model shows that these states can be maintained over very long timescales, and it allows to identify the mechanisms by which cohesion is lost in the long term.

## I FISH SCHOOL MODEL WITH BURST-AND-COAST SWIMMING

The “burst-and-coast” swimming mode consists of a successive alternation of sudden accelerations and quasi-passive deceleration periods during which the fish glides along a nearly straight line. These short events during which a fish changes both its velocity and its direction of motion are called “kicks”. Consecutive kicks of a single fish do not have necessarily the same length and duration. When swimming in groups, the kicks performed by different fish are mostly asynchronous and of different lengths and durations. We consider the duration of the acceleration phase to be much smaller than that of the gliding phase, and can therefore be neglected. We also consider that each fish chooses its direction of motion at the instant of performing a kick, and that it maintains its heading unchanged while decelerating along the kick.

We denote by 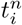 the instant of time at which fish *i* makes its *n*-th kick, and by 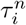 and 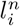 the duration and length of this kick, respectively. The position and velocity vectors of fish *i* at this kicking time are 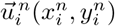 and 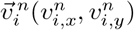 respectively.

The speed of the fish at the beginning of the kick is given by 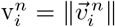. The speed is found experimentally to decay exponentially during the gliding phase with a fixed relaxation time *τ*_0_, while the fish heading orientation 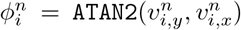 remains unchanged along the *n*-th kick. Positive angles are defined in the counter-clockwise direction, with respect to the positive semi-axis of abscissas of the global system of reference centered in (0, 0) (figure 1*a*, variables in red).

**FIG. 1.**
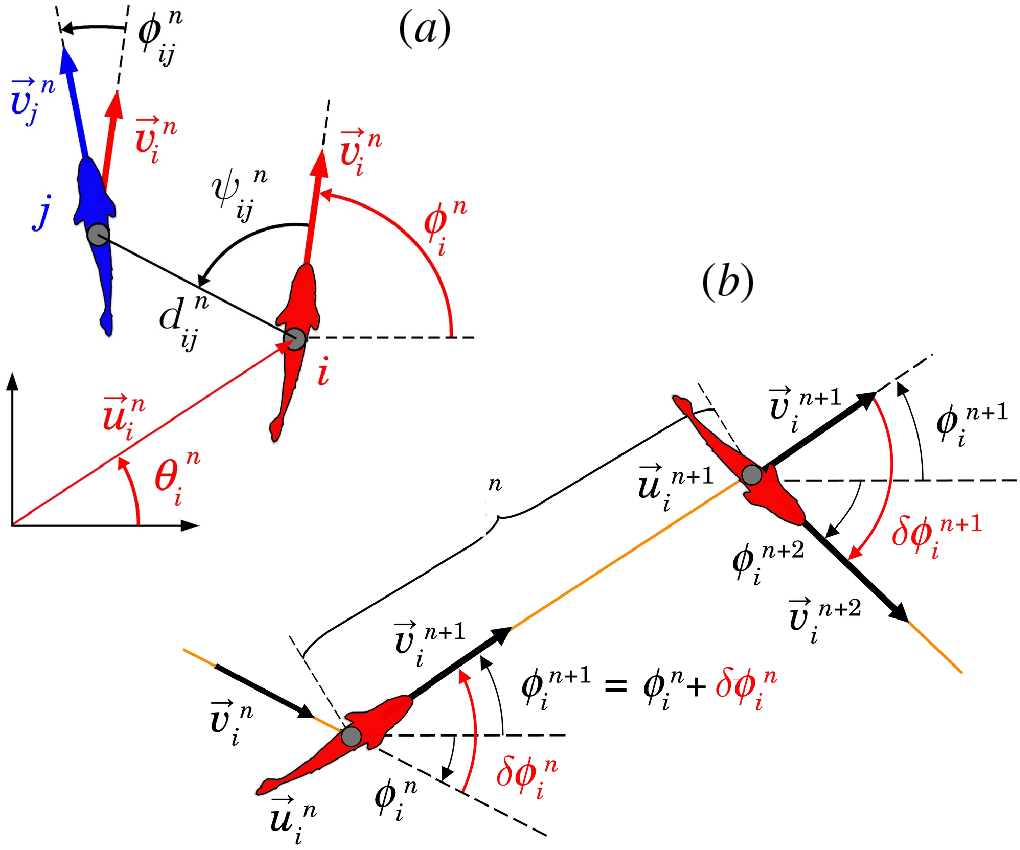
(a) Individual (red) and social (black) state variables (black) of fish *i* with respect to fish *j* at the instant 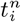 when fish *i* performs its *n*-th kick: 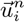 and 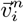 are the position and velocity vectors of fish *i*, 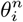 and 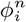 are the angles that these vectors make with the horizontal line, 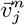 is the velocity vector of fish *j* at time 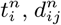 is the distance between fish *i* and fish *j* 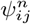 is the angle with which fish *j* is perceived by fish *i* (not necessarily equal to 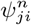), and 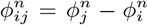 is the heading difference between both fish. (b) Schematic of the *n*-th kick performed by fish *i* moving from 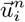 at time 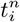 to 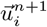 at time 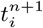 along a distance 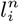. Orange lines denote fish trajectory, black wide arrows denote velocity vector, curved arrows rep-resent angles. The heading angle change of fish *i* at time 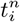 is 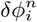. Fish heading during its *n*-th kick is 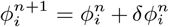. Red angles show the heading variation of the fish at the kick-ing instants 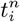 and 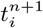.

The relative state of fish *i* and *j* is given by (*d*_*ij*_, *ψ*_*ij*_, *ϕ*_*ij*_), where *d*_*ij*_ is the distance between fish, *ψ*_*ij*_ = *θ*_*ij*_ −*ϕ*_*i*_ is the angle with which fish *i* perceives fish *j, θ*_*ij*_ is the angle that the vector going from *i* to *j* forms with the horizontal line, and *ϕ*_*ij*_ = *ϕ*_*j*_ − *ϕ*_*i*_ is the relative heading, which also measures how much *i* and *j* are aligned. Note that *ψ*_*ij*_ is not necessarily equal to *ψ*_*ji*_ (figure 1*a*, variables in black).

We generalize the model derived from experimental data in [13] and [18] by simplifying fish dynamics and social interaction functions while preserving the essential ingredients of the discrete and asynchronous burst-and-coast swimming mode and the interaction strategy of paying attention to only the one or two most influential neighbors. Social interactions are thus described by pairwise interaction functions of attraction and alignment, which scaled with the typical body length of fish, and where only the leading mode of the angular components is kept.

Figure 1*b* shows the *n*-th kick performed by fish *i* at time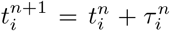. The position and heading of the fish at the end of its *n*-th kick is given by

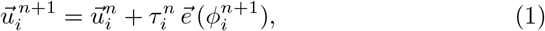

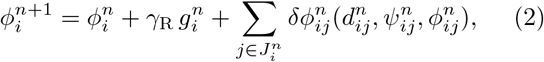

where 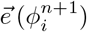 is the unitary vector pointing in the direction of the angle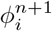.

The length and duration of the *n*-th kick of fish *i* are taken equal (unit speed scale), 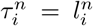, and sampled from a bell-shaped distribution of mean 1 (figure 2*a*).

**FIG. 2.**
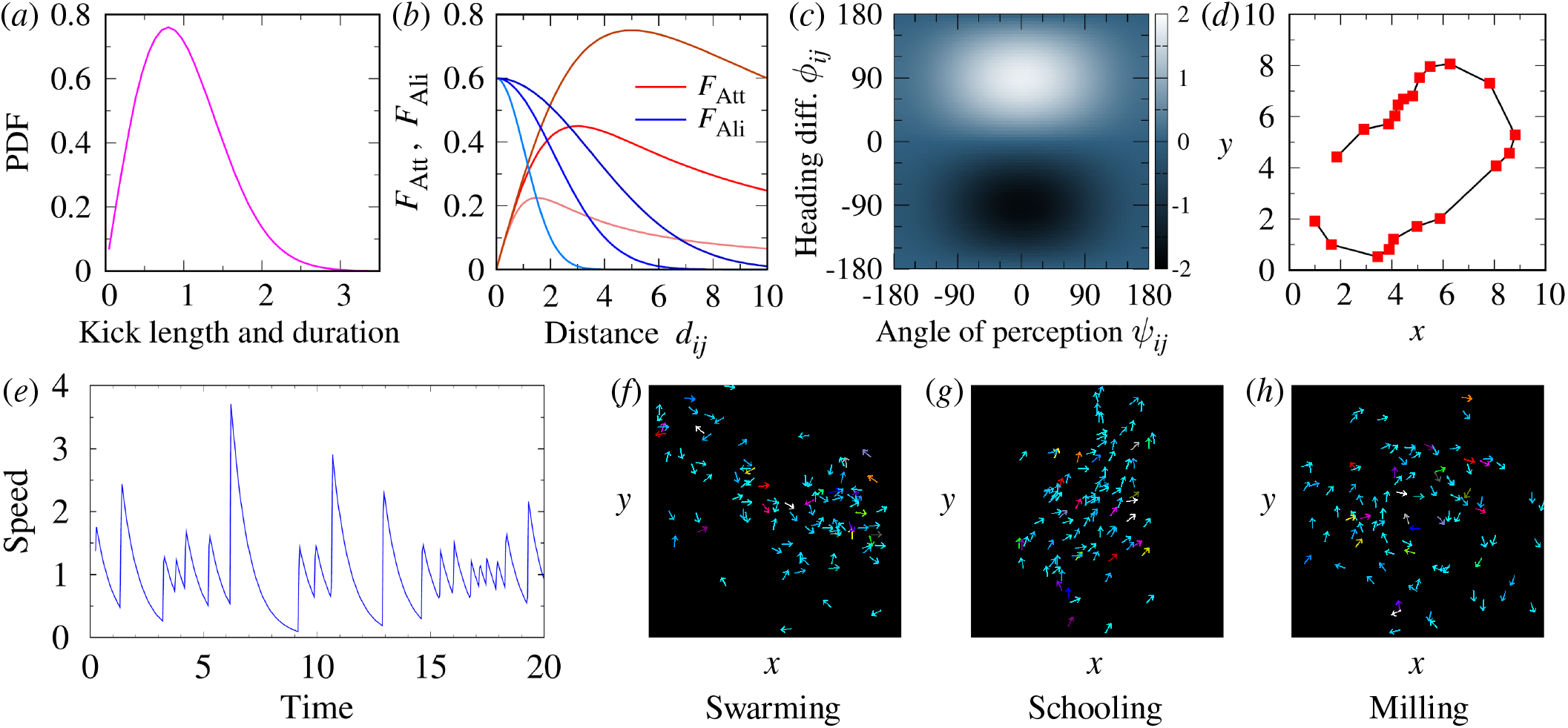
Individual burst-and-coast swimming behavior, social interaction between fish and collective phases observed in the model). (*a*) Probability density function (PDF) of the kick length and duration. (*b*) Social forces of attraction (red) and alignment (blue) as a function of the distance *d*_*ij*_ between fish for three values of the interaction ranges *l*_*Att*_ = *l*_*Ali*_ =1.5, 3 and 5. (*c*) Strength of alignment, as the product of an even function of the angle of perception *ψ*_*ij*_ and an odd function of the heading difference *ϕ*_*ij*_. (*d*) Trajectory of a single fish showing the kicking instants (red squares) followed by straight segments (black lines). (*e*) Velocity profile showing the gliding phases corresponding to the 20 kicks shown in (*d*). (*f–h)* Typical collective motion patterns of each phase: (*f*) swarming, (*g*) schooling, and (*h*) milling, for *k* = 1. Not all the *N* = 100 fish appear in the figures.

The heading angle change 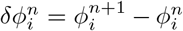 of fish *i* is the result of 1) the random fluctuations of the fish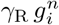, where *γ*_R_ is the fluctuation intensity and 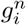 is a random number sampled from a Gaussian distribution of mean equal to one, and 2) the social interactions with other fish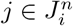, where 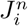 is the set of fish considered to have an influence on the behavior of fish *i*.

The *influence* that a fish *j* exerts on a fish *i* is precisely given by the absolute value of the heading angle change of fish *i* induced by fish *j* [18]:

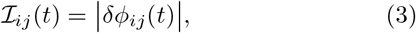

and therefore depends on the relative state of both fish. Sorting the neighbors of fish *i* according to ℐ_*ij*_(*t*) allows to identify the *k* = 1 or 2 most influential neighbors that will constitute the social environment according to the selected social interaction strategy. As shown in our recent results on *H. rhodostomus*, only the two most influential neighbors are required to quantitatively reproduce the behavioral patterns performed by groups of fish swimming in a circular tank [18]. As the influence that fish exert on a focal fish can change from one kick to another, the identity of the most influential neighbor(s) can change accordingly.

The strength of the social interaction depends non-linearly on the distance between fish (figure 2*b*), and is modulated by the relative position and heading of both fish (figure 2*c*). Under equal conditions of alignment, fish *i* will turn right if fish *j* is on its right side and will turn left otherwise, so that attraction is an odd function of *ψ*_*ij*_. Although experiments with pairs of *H. rhodostomus* have shown that attraction slightly depends on the way fish are aligned with each other, we assume here that the relative heading *ϕ*_*ij*_ has no effect on the attraction strength. Similarly, alignment is an odd function of *ϕ*_*ij*_: for fish j located at a given place, fish *i* will turn right if *ϕ*_*ij*_ > 0, and will turn left otherwise. Alignment also depends on the relative position of the perceived fish, but in an even way: the alignment force exerted by a neighbor located in front of a fish (|*ψ*_*ij*_ | < *π*/2) is stronger than when the neighbor is located behind (| *ψ*_*ij*_ |> *π*/2), but does not depend on whether the neighbor is located on the right (*ψ*_*ij*_ < 0) or on the left side (*ψ*_*ij*_ > 0) of the focal fish.

Assuming, as in [13], that attraction and alignment are combined in an additive form, and that all contributions can be decoupled in functions of a single state variable, the heading angle change of fish *i* due to social interaction with one neighbor *j* is given by

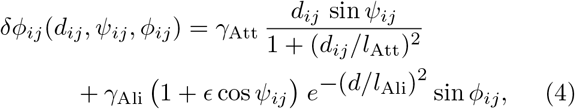

where all parameters are dimensionless: *γ*_Att_ and *γ*_Ali_ are the respective intensities of the attraction and alignment forces, *l*_Att_ and *l*_Ali_ the respective interaction ranges (scaled with the typical kick length *l*_0_ = 0.07 m), and ϵ is a parameter of anisotropy in the alignment. The first term in Eq. (4) corresponds to the attraction, and the second to the alignment. Both terms have been simplified with respect to the original data-based model [27], by removing the short range repulsion and alignment, and retaining only one Fourier mode in the angular components.

Figure 2*b* shows the strength of attraction and alignment as a function of the distance between fish for three values of the respective range of interaction *l*_Att_ and *l*_Ali_. The modulation of these forces as functions of the angle of perception and the degree of alignment is a simple sin *ψ*_*ij*_ for the attraction and a two-dimensional function of both angles for the alignment (figure 2*c*).

In order to calculate *δϕ*_*ij*_, the relative state of fish *j* must be known at the kicking time of fish *i*. As kicks of different fish are asynchronous, the position and heading angle of fish *j* must be calculated according to the elapsed time since its last own kick. The position of a fish *i* during the gliding phase following its *n*-th kick, *i*.*e*., at time 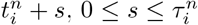 can be written as

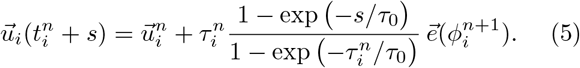

These intermediate positions also allow to represent the trajectories of the *N* fish with a fixed time step.

Figures 2*d* and 2*e* show respectively the trajectory of a single fish along 20 kicks and the corresponding speed profile, both illustrating the discrete nature of the burst-and-coast swimming mode. Despite the fact that fish collect information from the environment only at kicking instants, that their environment is limited to one or two specific neighbors, and that their decision is locked along their kicks, groups of burst-and-coasting fish are able to display highly structured collective behaviors. Figures 2*f–h* show the result of a single run of the model in a large group of *N* = 100 individuals and for three pairs of values of the social interaction parameters, (*γ*_Att_, *γ*_Ali_) = (0.6, 0.6), (0.22, 0.6), and (0.37, 0.2), which give rise to a cohesive school displaying respectively a swarming, a schooling, and a milling formation.

## II COLLECTIVE MOTION PATTERNS AND PHASE DIAGRAM

We first explore the parameter space of the model applied to a group of *N* = 100 fish by varying its main parameters, which are the intensity of attraction and alignment, *γ*_Att_ and *γ*_Ali_, respectively.

Figure 3 shows the dispersion, polarization, and milling, obtained for different values of the intensity of the attraction and alignment *γ*_Att_ and *γ*_Ali_ while keeping other parameters fixed and for both social interaction strategies, *k* = 1 and *k* = 2.

**FIG. 3.**
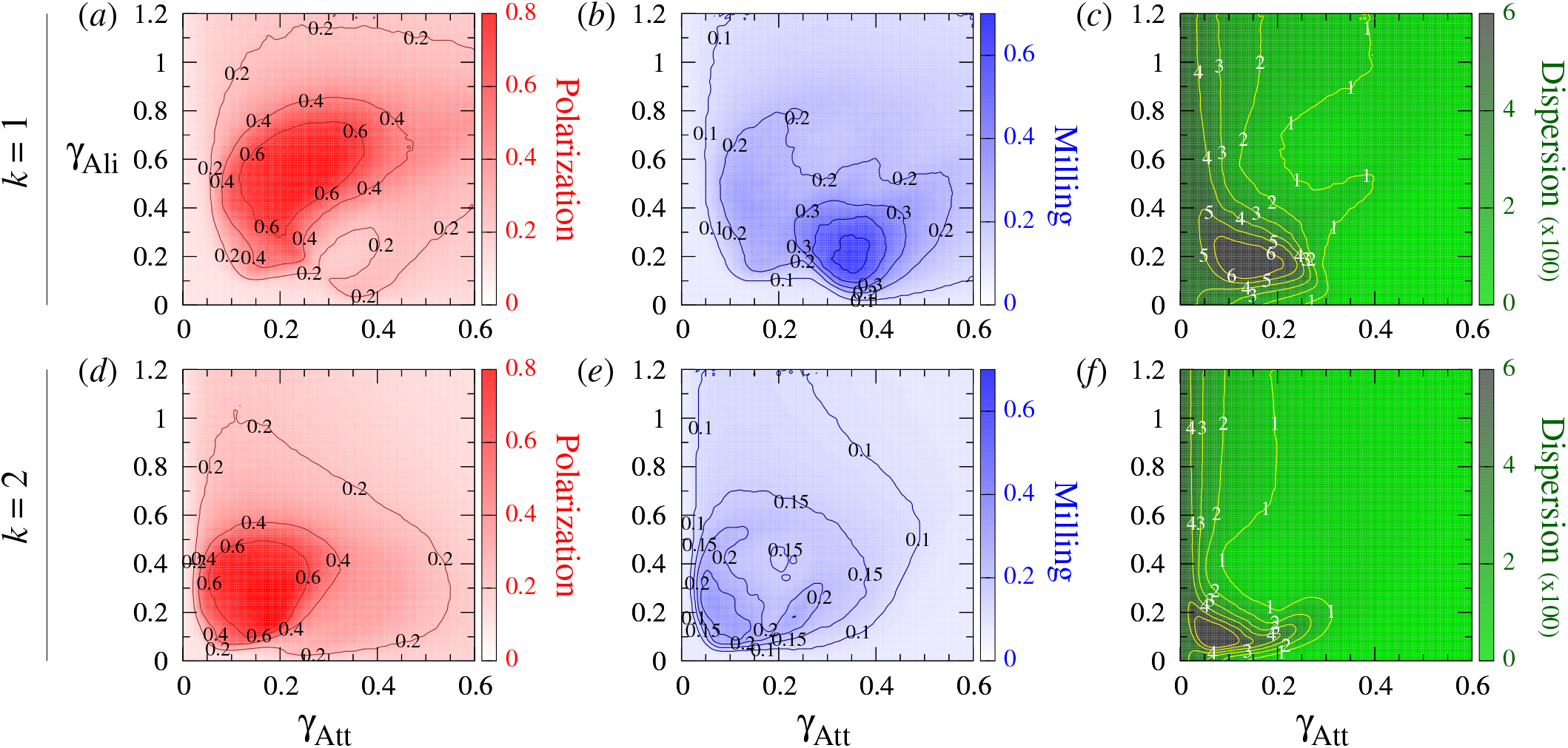
Polarization, milling, and dispersion maps for different values of the intensity of attraction and alignment strengths *γ*_Att_ and *γ*_Ali_ respectively, when fish interact with their most influential neighbor (*k* = 1) and their two most influential neighbors (*k* = 2). (*ad*) Polarization (red). (*be*) Milling (blue). (*cf*) Dispersion (green). Regions of high color intensity mean that fish frequently display the characteristic behavioral patterns of schooling, milling, or swarming. The green region means that the school of fish is highly cohesive. In the gray region, attraction is too weak and fish quickly disperse. Social interaction ranges are *l*_Att_ = *l*_Ali_ = 3m, ϵ = 0.8, intensity of random fluctuation (noise) is *γ*_R_ = 0.2. Each pixel is the average of 20 runs of 2000 kicks per fish (the first half has been discarded in order to remove the effects of the initial condition). The maps have been smoothed to enhance the representation of level sets.

When *k* = 1 (figures 3*a–c*), dispersion is uniformly small (*D* < 150) in the right half plane where the attraction strength is large (*γ*_Att_ > 0.3). For smaller values of *γ*_Att_, dispersion is higher and, depending on the alignment strength, reaches very high values (*D* > 600) in a small region where *γ*_Att_ ∈ (0.05, 0.25) and *γ*_Ali_ ∈ (0.1, 0.3). Polarization (figure 3*b*) reaches high values (*P* > 0.4 and up to 0.8) in a large kidney-shaped region where *γ*_Att_ ∈ (0.07, 0.4) and *γ*_Att_ ∈ (0.1, 0.8). Outside this region, the group is not polarized, *P* ≈ 0.1–0.2. Indeed, note that for *N* = 100 individuals, the expected value of the polarization corresponding to uncorrelated headings is of order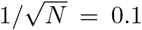. Similarly, there is a region of high milling where *M* > 0.3 (up to 0.7 in the center), located around (*γ*_Att_, *γ*_Ali_) = (0.35, 0.2), and whose size is about one third of the region of high polarization. Outside this region, the milling is uniform and small (*M* < 0.1). We also observe a slightly visible region of intermediates values of milling (*M* **≈** 0.2) whose boundary coincides with that of the region of high polarization.

For *k* = 2 (figures 3*d–f*), there is no region of high milling. The regions of high polarization and high dispersion are smaller than for *k* = 1, and are obtained for smaller values of the interaction strengths *γ*_Att_ and *γ*_Ali_.

Figure 4*a* shows the collective behavioral phases for the interaction strategy *k* = 1, obtained by superimposing these regions of high values in a phase diagram. We define the following three phases of collective behavior:

**FIG. 4.**
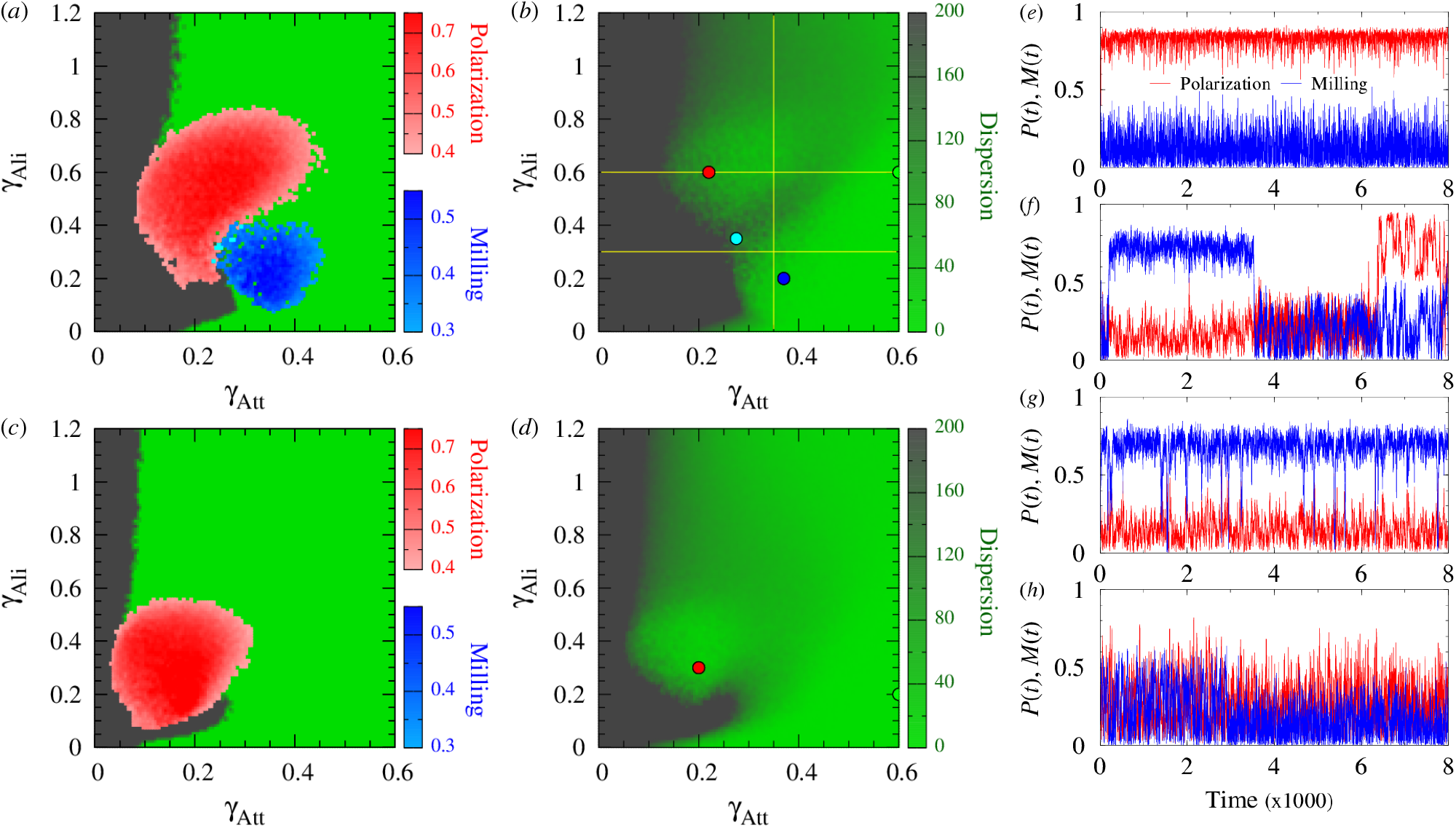
Phase diagrams *γ*_Att_-*γ*_Ali_ showing the regions of high polarization (*P* ≥ 0.4) and high milling (*M* ≥ 0.3) determining the phases of schooling (red) and milling (blue) respectively, for (*a*) *k* = 1 and (*c*) *k* = 2, superimposed to a binary color map of dispersion delimiting the swarming phase (*D* ≤ 200, light green), where the group stays cohesive, and a high dispersion region (*D* > 200, gray). Points where *P* ≥ 0.4 and *M* ≥ 0.3 simultaneously are shown in cyan. (*b*) Color map of dispersion for *D* ∈ [0, 200] and *k* = 1, and (*d*) for *k* = 2, showing the three transects (yellow lines) at *γ*_Att_ = 0.35, *γ*_Ali_ = 0.6, and *γ*_Ali_ = 0.3, detailed in figure 5. In (*b*) and (*d*), values larger than *D* = 200 appear in dark green. The four colored points in (*b*) are at (0.22, 0.6) (red), (0.37, 0.2) (milling), (0.275, 0.35) (cyan), and (0.6, 0.6) (green). The two points in (*d*) are at (0.2, 0.3) (red) and (0.6, 0.2) (green). (*e–h*) Time series of polarization *P*(*t*) (red lines) and milling *M*(*t*) (blue lines) corresponding to the points (*γ*_Att_, *γ*_Ali_) shown in panel (*b*): (*e*) red point, high polarization and low milling; (*f*) cyan point, alternating high and low values of *P*(*t*) and *M*(*t*); (*g*) blue point, high milling and low polarization; (*h*) green point, small values of P(t) and M(t). Other parameters are *γ*_R_ = 0.2, *l*_Att_ = *l*_Ali_ = 3, and ϵ = 0.8. Each point is the average of 20 runs of 2 × 10^4^ kicks per fish (first half discarded in order to remove the effects of the initial condition).

- Schooling (red): *P* ≥ 0.4 and *M* < 0.3;
- Milling (blue): *P* < 0.4 and *M* ≥ 0.3;
- Swarming (green): *P* < 0.4 and *M* < 0.3.

We show in cyan the points where *P* ≥ 0.4 and *M* ≥ 0.3.

Figure 4*b* shows the dispersion with two uniformly colored regions of low (green) and high (gray) values for *k* = 1. When the dispersion is small, the group remains cohesive at least for the duration of the simulations. The region of high dispersion cannot be considered a collective phase. For *k* = 1 (figures 4*a,b*, see also electronic supplementary material, video S1, video S2, video S3 and video S4), a minimal attraction strength 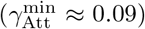 is necessary to prevent dispersion. The polarization (red) and milling (blue) phases are mainly located within the low dispersion region, but they also cover partly the region of high dispersion. The polarization and milling phases are located next to each other, with a very small region inbetween (cyan) in which fish alternate between schooling and milling (figure 4*f*). For higher values of the attraction and alignment strengths, there is no more schooling or milling and the fish adopt a swarming behavior (green).

For *k* = 2 (figures 4*c,d*, see also electronic supplementary material, video S5 and video S6), we first observe that there is no milling phase, meaning that *M* < 0.3 for all *γ*_Att_ and *γ*_Ali_. The minimal attraction strength to prevent dispersion is smaller than when *k* = 1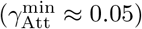, and the schooling phase is smaller than when *k* = 1. Fish start schooling at considerably smaller values of *γ*_Att_ (less than half the value leading to schooling when *k* = 1) and *γ*_Ali_ (less than 2/3 the corresponding value when *k* = 1), and reach higher polarization values. No schooling appears when *γ*_Att_ > 0.3 or *γ*_Ali_ > 0.55. Similarly, when *γ*_Ali_ > 0.2, the swarming phase starts at smaller values of *γ*_Att_ compared to the condition with *k* = 1. The transition from high to low dispersion is located at half the value of *γ*_Att_ observed for *k* = 1. In turn, for smaller values of *γ*_Ali_, the region of dispersion is similar to the one observed when *k* = 1, now forming a tongue that enters into the swarming region up to *γ*_Att_ ≈ 0.3. For very small *γ*_Ali_ < 0.05, the swarming phase starts at values of *γ*_Att_ that are half those found for *k* = 1 (0.1 instead of 0.2).

Figures 4*e–h* show the time series of polarization and milling for the four points plotted in the phase diagram in figure 4*b* (see also electronic supplementary material, video S1, video S2, video S3 and video S4). For *γ*_Att_ and *γ*_Ali_ picked in the center of the schooling region, polarization remains almost all the time above *P* = 0.8 (figure 4*e*). The same happens for *M*(*t*) when (*γ*_Att_, *γ*_Ali_) is in the center of the milling phase (figure 4*g*), reaching and maintaining a stable value around *M* = 0.75, although the milling structure is sometimes lost (but quickly recovered) in this example. At the interface of the schooling and milling regions (figure 4*f*), both order parameters are significant, on average. This results from an intermittent dynamics where the school can switch back and forth from a polarized to a milling organization. This interfacial region corresponds to large fluctuations of both order parameters, and also to a strong sensitivity of the group to any external perturbation [28]. Figure 4*h* illustrates the case where (*γ*_Att_, *γ*_Ali_) is in the swarming phase; even in that case, the group adopts quite often a spatial configuration that gives rise to non-negligible values of the polarization, up to *P* = 0.6, but only for very short intervals of time. The same happens with milling (*M*(*t*) is often above 0.4).

In order to quantify the extent to which the instantaneous polarization and milling are larger than their respective mean, we calculate the probability density function (PDF) of *P*(*t*) and *M*(*t*) along three transects that cross the behavioral phases of the phase plane (yellow lines in figures 4*b*).

Figure 5 shows that, across the behavioral phases of schooling and milling, the corresponding PDF typically exhibits a peak at high values and a long tail, precisely responsible for the low mean value. The peak of the PDF of polarization is at about *P* = 0.8 and *P* = 0.85 in the two transects crossing the schooling phase, while the mean is at about *P* = 0.65 and *P* = 0.73 respectively (figure. 5*a,c*). Similarly, the peak of the milling is at about *M* = 0.7, while the mean is always smaller than *M* = 0.6 (figure 5*b,f*).

**FIG. 5.**
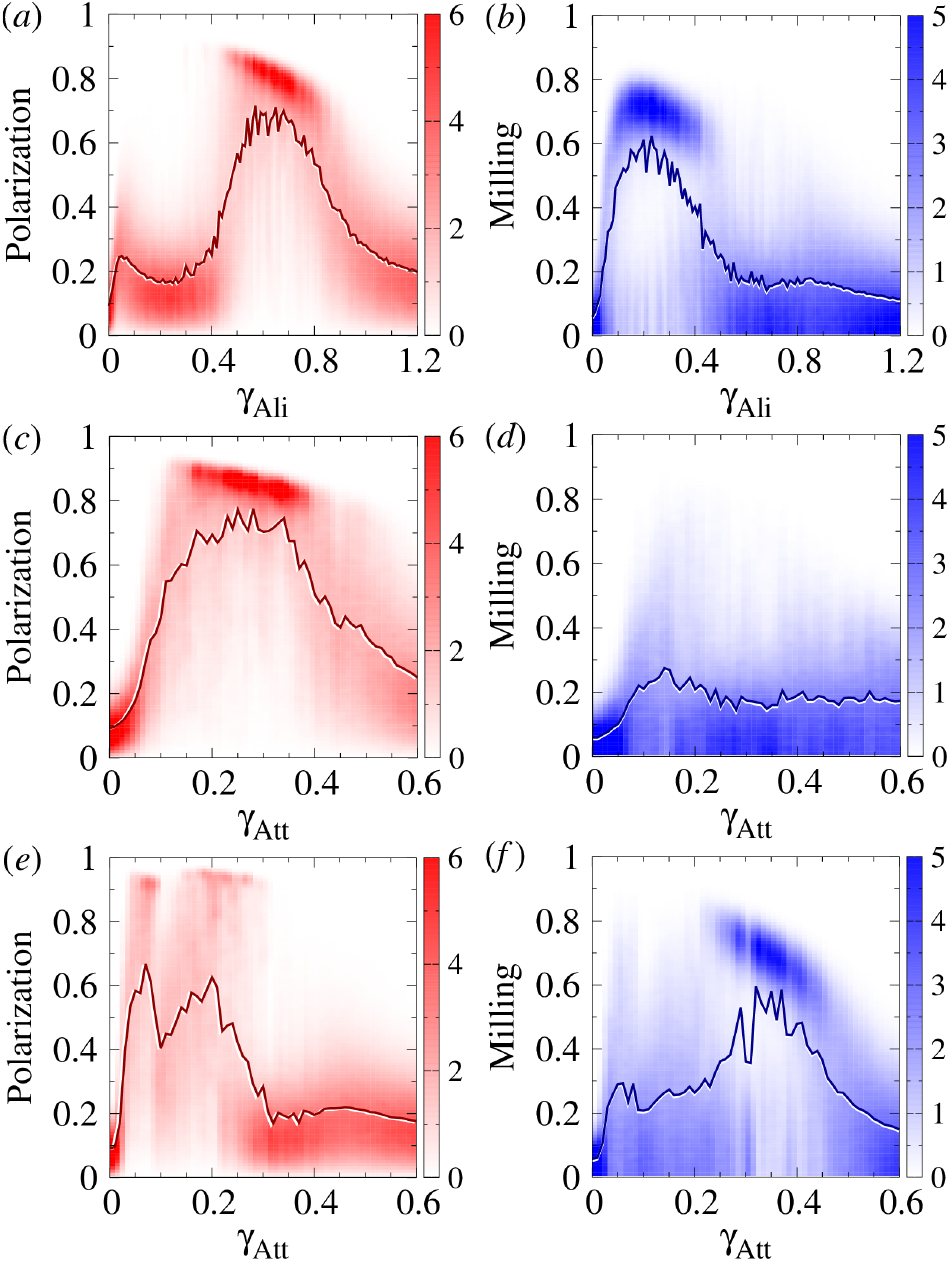
Polarization (red) and milling (blue) probability distribution functions (PDF) and mean values (solid lines) along the transects of the phase diagram for *k* = 1 (yellow lines in Fig. 4*b*). (*a,b*) Vertical transect at *γ*_Att_ = 0.35 crossing the three phases, (*c,d*) horizontal transect at *γ*_Ali_ = 0.6 crossing the polarization and swarming phases, and (*e,f*) horizontal transect at *γ*_Ali_ = 0.3 crossing the milling and swarming phases. In all panels, the variation of color intensity along the *y*-axis corresponds to the PDF of the parameter value shown in the *x*-axis, and solid lines join the corresponding mean value of each PDF. The rest of parameters are *γ*_R_ = 0.2, *l*_Att_ = *l*_Ali_ = 3, ϵ = 0.8, and *γ*_R_ = 0.2.

Outside the schooling and milling phases, both polarization and milling are widely and uniformly distributed around the mean value; see, *e*.*g*., *γ*_Ali_ ∈ [0, 0.4] in figure 5*a* and *γ*_Att_ ∈ [0.3, 0.6] in figure 5*e* for polarization, and *γ*_Ali_ ∈ [0.5, 1.2] in figure 5*b* and the whole transect in figure 5*d* for milling. Moreover, the higher the dispersion, the greater the width of the PDF. For *γ*_Ali_ ∈ [0, 0.4] in figure 5*a*, the dispersion is quite small (*D* < 40) and the PDF of polarization is quite narrow, in comparison with the interval *γ*_Att_ ∈ [0.4, 0.5] in figure 5*c*, where the dispersion is higher (*D* ≈ 80) and the PDF wider, and with the interval *γ*_Att_ ∈ [0.025, 0.275] in figure 5*e*, where the dispersion is very high (*D* > 200), and the PDF is practically flat.

The same happens for milling, comparing the interval *γ*_Ali_ ∈ [0.6, 1.2] in figure 5*b*, where the dispersion is low and the PDF is narrow, with the interval *γ*_Att_ ∈ [0.025, 0.275] in figure 5*f*, where the dispersion is very high and the PDF is almost flat.

## III LONG-TERM DYNAMICS AND STABILITY OF COLLECTIVE PHASES

Schooling and milling are complex dynamic structures that can be destabilized by different events such as the sudden change of behavior of a subgroup of individuals (leading to the splitting of the fish school), the simul-taneous departure of small subgroups of fish (leading to the scattering of the fish school), or the distancing of a few individuals who lose contact with the school and gradually move away from the group (producing a slow disintegration of the fish school).

In these three cases, very long simulations must be carried out to study the evolution of the dispersion and determine how the stability of the spatial structures is lost. We thus characterize the stability by monitoring the dispersion of the fish school and the number and size of subgroups, along a large amount of long simulations (100 runs of 2 × 10^4^ kicks per fish), keeping track of the state of the *N* = 100 individuals every 20 kicks, for *γ*_Ali_ = 0.6 and *γ*_Att_ ∈ [0, 0.6] and *k* = 1, which is the upper horizontal transect of the phase plane shown in figure 4*b*.

We used a discretization step Δ*γ*_Att_ = 0.025, attraction and alignment ranges *l*_Att_ = *l*_Ali_ = 3, *ϵ* = 0.8, *γ*_R_ = 0.2, and an initial condition where individuals’ position and heading are sampled from a uniform random distribution.

Figure 6 shows the time evolution of the mean polarization and the mean squared dispersion *D*^2^(*t*). As expected, the mean polarization reaches its highest values in the schooling phase, when *γ*_Att_ ∈ (0.15, 0.375) (red lines). Outside this region, the mean polarization of the school is much smaller for two different reasons, depending on whether *γ*_Att_ lies in the dispersion region (*γ*_Att_ < 0.15) or in the swarming phase (*γ*_Att_ > 0.375). When *γ*_Att_ < 0.15, the mean squared dispersion quickly grows with time and reaches very high values (dark red lines in figure 6*b*). At the same time, the mean polarization decreases to the value of random polarization (*P*_0_ = 0.1). Furthermore, the smaller *γ*_Att_, the faster the changes in polarization and dispersion. In turn, when *γ*_Att_ > 0.375 (green lines), the squared dispersion is about 3 decades smaller than when *γ*_Att_ < 0.15, and no decay in time is observed in the average polarization.

**Figure 6.**
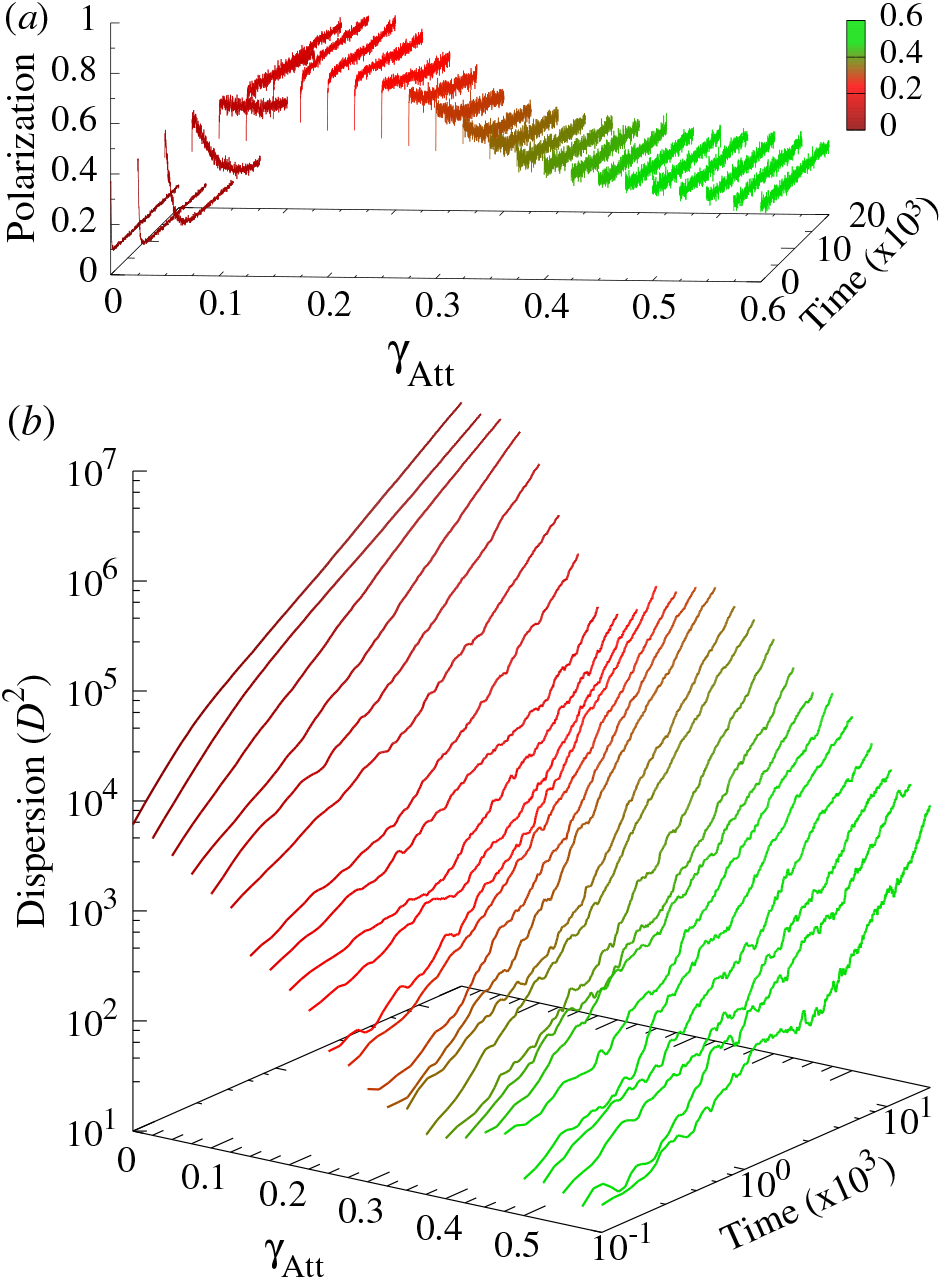
Time series of (*a*) mean polarization *P*(*t*) and (*b*) mean squared dispersion *D* ^2^(*t*) averaged over 100 long simulations (2 × 10^4^ kicks per fish) for *γ*_Ali_ = 0.6 and *γ*_Att_ ∈ [0, 0.6] and *k* = 1 (corresponding to the upper horizontal cut of the phase plane in figure 4*b*). Time and dispersion scales are logarithmic in (*b*). The colors correspond to those used in figure 4*a* to represent dispersal region and the schooling and swarming phases. Other parameter values: *l*_Att_ =*l*_Ali_ = 3, *ϵ* = 0.8, *γ*_R_ = 0.2.

This suggests that, for small *γ*_Att_, polarization is small because the group quickly fragments and the alignment is not preserved between distant subgroups, even if the alignment is strong enough to allow individuals to align with each other inside each subgroup. On the other hand, for large values of *γ*_Att_, polarization is small because individuals fail to align with each other despite the fact that the group remains cohesive. Indeed, when the attraction becomes too strong, the heading change of the focal fish resulting from the interaction with its most influential neighbor becomes too large and effectively random, leading to a decrease in the group polarization.

To check this, we calculate the time evolution of *i*) the mean dispersion *D*(*t*) of the *N* = 100 fish, *ii*) the number of subgroups *N*_G_(*t*), *iii*) the size of the largest subgroup, and *iv*) the mean dispersion inside subgroups *D*_G_(*t*), for values of (*γ*_Att_, *γ*_Ali_) corresponding to the points shown in each phase of the phase planes of figures 4*b,d* for both *k* = 1 and *k* = 2. We used a much larger number of 1000 even longer runs of 4×10^4^kicks per fish. Moreover, as the initial random heading of the fish can favor the fragmentation of the group, we take an initial condition where all individuals have the same heading and are positioned in an elliptic spatial configuration with an inter-distance similar to the one observed when the group is schooling. The number of groups is calculated with a recursive algorithm where the group to which a fish belongs contains its nearest neighbor, the nearest neighbor of its nearest neighbor, and so on, and a critical distance is used to consider that close groups are, in fact, the same group; see Sec. 5c.

Figure 7a shows that, for *k* = 1, dispersion grows in the three phases of schooling, milling, and swarming. However, the growth rate is much higher in the schooling and milling phases than in the swarming phase. After *t* ≈ 5000, dispersion in the milling phase is always half a decade above the one observed in the schooling phase, and two decades above the one observed in the swarming state at the end of the simulations (*t* = 4×10^4^). Dispersion in the schooling state is one decade and a half higher than the one observed in the swarming state. The time evolution of the mean number of groups over these 100 runs shows that the high dispersion in the schooling and milling phases is due to the fragmentation of the group. Figures 7*a,b* show that the growth rate of the dispersion in schooling (red) and milling (blue) phases increases precisely when the corresponding number of groups departs from *N*_G_ = 1, at *t* ≈ 3000 in the schooling case, and sooner, at *t* ≈ 1000 in the milling phase. In turn, the group remains cohesive in the swarming phase (*N*_G_ ≈ 1 until the final state).

**Figure 7.**
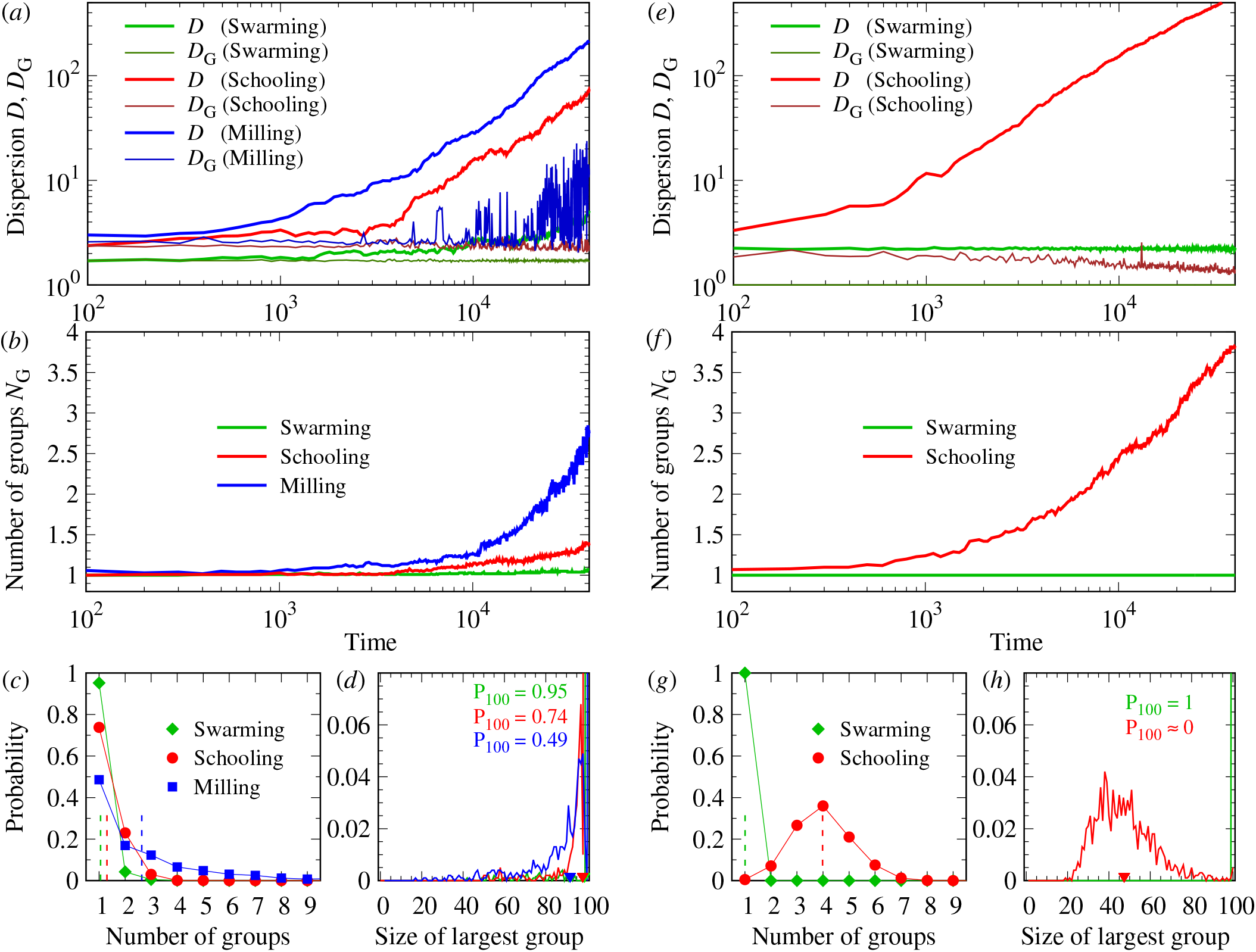
Dispersion, number of groups, and size of the largest subgroup, for the three points (*γ*_Att_, *γ*_Ali_) in the swarming (green), schooling (red), and milling (blue) phases shown in figures 4*b,d*, for *k* = 1 (*a–d*) and *k* = 2 (*e–h*). (*a,e*): Time evolution of the mean dispersion of the whole group *D*(*t*) and of the mean dispersion averaged over all subgroups *D*_G_(*t*) (log-log scale). (*b,f*): Time evolution of the mean number of groups *N*_G_(*t*) (log-normal scale), corresponding to (*a*) and (*e*) respectively. Each line is the average over 1000 runs (4 × 10^6^ kicks per run). (*c,g*): Probability distribution (symbols) and mean value (dashed lines) of the number of groups *N*_G_, calculated on the final state of each of the 1000 runs. (*d,h*): Probability distribution (solid lines) and mean value (small triangles on the *x*-axis) of the size of the largest group. The curves of schooling and milling are shifted ™1 and +1 respectively not to overlap the peaks of probability at 100. The range of abscissas is thus extended to [™1, 101]. Colored numbers denote the height of each peak. Parameter values when *k* = 1 are (see the points in figure 4*b*): swarming: (*γ*_Att_, *γ*_Ali_) = (0.6, 0.6); schooling: (*γ*_Att_, *γ*_Ali_) = (0.22, 0.6); milling: (*γ*_Att_, *γ*_Ali_) = (0.37, 0.2). Parameter values when *k* = 2 are (see the points in figure 4*d*): swarming: (*γ*_Att_, *γ*_Ali_) = (0.6, 0.2); schooling: (*γ*_Att_, *γ*_Ali_) = (0.2, 0.3). Initially fish have the same heading (*ϕ*= 0) and are distributed in an ellipse of long and short half axes *a* and *b* respectively (long axis taken parallel to the *x*-axis). See the rest of parameter values in Table 1.

Once separated from the main group, subgroups behave differently in the schooling and the milling phases. Figure 7*a* shows that, in the schooling phase, the mean dispersion inside subgroups remains constant in time (*D*_G_(*t*) ≈ 2.5, dark red line in figure 7*a*) and the mean number of subgroups at the final state is very small (*N*_G_ < 1.5, red line in figure 7*b*). However, in the milling case, the mean dispersion inside subgroups continues to grow, especially after *t* ≈ 10000 (*D*_G_ = 10 at the final state, dark blue line in figure 7*a*), thus inducing new separations and increasing the number of subgroups *N*_G_ until the final time (blue line in figure 7*b*).

**TABLE 1.** Parameter values used in the simulations. (*a*) Interaction strength and range, noise, mean kick duration and length, relaxation time, and coefficient of anisotropy; (*b*) long and short axes a and b repectively of the ellipse inside which fish are randomly distributed in the initial conditions.

| (a) |  | (b) |  |  |
| --- | --- | --- | --- | --- |
| Parameter | Values | $k$ | Phase | $a$ $b$ |
| $\gamma_{\text{Att}}$ | 0–0.6 | $k=1$ | swarming | 3 1.5 |
| $\gamma_{\text{Ali}}$ | 0–1.2 | | schooling | 5 3 |
| $l_{\text{Att}}$ | 1.5, 3, 5 | | milling | 3 1.5 |
| $l_{\text{Ali}}$ | 1.5, 3, 5 | $k=2$ | swarming | 4 2 |
| $\gamma_{\text{R}}$ | 0.2 | | schooling | 3 1.5 |
| $\langle \tau_n \rangle, \langle l_n \rangle$ | 1 | | milling | – – |
| $\tau_0$ | 0.8 | | | |
| $\varepsilon$ | 0.8 | | | |

Figures 7*c,d* show the probability of having a given number of groups in the final configuration, and the probability that the largest group has a given size at the final configuration, calculated over the 1000 simulations. The fraction of final states where only one group is present is 95% in the swarming phase, 74% in the schooling case, and 49% in the milling phase. Configurations with 3 groups are never found in the swarming phase, and quite rarely in the schooling and milling phase. In all phases, the largest group is very large (mean size larger than 90, figure 7*d*). This means that the typical configuration consists of a single very large group and very few small groups, with no groups of intermediate size. Separation almost never happens in the swarming phase, and is less frequent in the schooling phase than in the milling case, where more subgroups are found.

For *k* = 2, we only consider two points of the phase plane, as there is no milling phase (figure 4*d*). In the swarming phase, the mean dispersion remains at the same value during the whole simulation time (figure 7*e*), and there is only a single group in all final states (figure 7*f*).

In turn, in the schooling case, the dispersion and the fragmentation are much higher than for *k* = 1. The growth rate of the mean dispersion starts to increase much sooner than for *k* = 1, at *t* ≈ 1000 instead of 4000, and at a similar rate, so that the curve of *D*(*t*) is almost half a decade above that observed for *k* = 1 during the second half of the simulations.

At the final time, *D* ≈ 548 when *k* = 2 while *D* ≈ 71 when *k* = 1. The mean number of groups at the final time is also higher than for *k* = 1, with a peak at *N*_G_ = 4 (figure 7*g*). However, the mean dispersion inside the subgroups remains constant and even slightly decreases (figure 7e, dark-red curve).

The group remained cohesive only in 5 of the 1000 runs. Final states with 5 groups or more are not rare. Moreover, the largest group is now of moderate size (*G*_1_ ≈ 35–55, with a mean value *G*_1_ ≈ 47.39, figure 7h). This means that, when *k* = 2, dispersion is so fast that the subgroups that separate from the main one are not necessarily small, so that a continuous range of sizes in 20–80 exists at the final state for the largest group, and consequently, a continuous range of small sizes for the smaller subgroups.

## IV DISCUSSION

We have introduced a simplified asynchronous burst-and-coast fish school model to investigate the impact of social interactions and interaction strategies between fish on the emerging collective phases and their long-term stability. The dynamics of the intermittent and asynchronous swimming mode is essentially discrete in time and space, and this property has a deep impact on the resulting collective states at the group level. Previous studies have shown that the typical gliding distance is one of the dominating scales that determine the emergence of group polarization [29], and that intermittent swimming modulates the influence of social interactions between fish on collective states [27]. These properties are tightly related to the unique way fish process social information during intermittent swimming, being only sensitive to that information just before the onset of the bursting phase while neglecting it during the gliding phase [13, 30]. This type of information processing contrasts with the one classically considered in most fish school models, in which each individual has a smooth speed and continuously updates its direction according to the direction and relative position of the other fish in a defined neighborhood [31–36].

This simplified model is able to reproduce the main features of the original data-based model [27]. First, the model shows that social interactions must reach a minimum intensity to keep a compact school and allow the emergence of coordinated states such as schooling and milling. However, stronger social interactions between fish do not necessarily lead to a higher level of coordination at the group level, and excessively strong social interactions can even prevent coordination. As captured by the vertical transect shown on figure 4*b*, for high values of alignment strength, the school entirely loses its polarization, and its dispersion also slightly increases. This phenomenon does not appear in models with continuous swimming dynamics [9, 36], in which the cohesion and the polarization of the school increase when the strength of social interactions becomes higher. As a matter of fact, when a fish chooses a new heading before performing a kick, high values of *γ*_Att_ and *γ*_Ali_ generate large *δϕ* in Eq. (2). The variation of headings between fish then increases, ultimately leading to a loss of coherence in their collective movements. When *δϕ* is even larger, fish behave like random walkers, and the group gradually disperses.

The model also reproduces the fact that when fish only interact with their most influential neighbor (*k* = 1), the region of the parameter space corresponding to the schooling and milling states is located within a narrow range of values of *γ*_Att_ and *γ*_Ali_, which also overlaps the transition region between group dispersal and the swarming state, meaning that collective states are sensitive to small variations of the social interactions’ strength.

Moreover, an important result of the model lies in the existence of a bi-stable region in which the fish school alternates between schooling and milling, at the boundary between these two states. Previous studies on species with an intermittent swimming mode, such as rummy-nose tetra (*H. rhodostomus*), zebrafish (*D. rerio*), and golden shiners (*N. crysoleucas*), have shown that, when swimming in group, fish regularly alternate between schooling and milling states [37–39]. As a matter of fact, operating near the transition region between collective states enhances information processing and adaptability in response to environmental cues [28, 40–42]. All these works suggest that these species have the capacity to collectively tune the strength of social interactions (governed by *γ*_Att_ and *γ*_Ali_ in the model) such that the system operates at a critical state, as it has recently been shown in shoals of sulfur molly (*Poecilia sulphuraria*) in natural conditions [37, 43].

Finally, the model is also able to show that the interaction with the two most influential neighbors allows the emergence of coordinated states with lower values of the alignment and attraction forces. When *k* = 2, the minimum values of *γ*_Att_ and *γ*_Ali_ leading to the schooling state are ∼ 2/3 of those found when *k* = 1. However, the region of the phase diagram leading to schooling state is much smaller than when *k* = 1 and the region in which milling was found is entirely replaced by swarming. This suggests that interaction of fish with their two most influential neighbors results in a much stronger effect of social interactions and a reduction in the parameter space, leading to ordered states.

Once the model is validated, we investigate the long-term stability of collective states. The first observation is that these collective states are quite stable, in the sense that they are sustained along a time that by far exceeds the time during which environmental conditions can reasonably be expected to remain unchanged in nature. However, for much longer times, the cohesion is lost for almost all cases.

When fish interact only with their most influential neighbor (*k* = 1), the fish school loses overall polarization and milling due to the formation of subgroups. The typical configuration is then one major group (of size ∼ 90 fish) which coexists with several small groups (figure 7*a–d*). In the three states of schooling, milling, and swarming, subgroups continue to disperse over the long term (figure 7*ab*), although dispersion starts before in the milling state, and is much slower in the swarming state. When fish interact with their two most influential neighbors (*k* = 2), the stability of the schooling state is reduced and there is no milling state. By contrast, the swarming state remains cohesive. This is a consequence of the stronger social influence experienced by fish.

Note that the emergence of subgroups resulting in a maximal typical size for cohesive groups can be biologically relevant for a given species. On the other hand, in our model based on the interaction with a very few most influential neighbors, larger groups could be stabilized by having individuals also interact with the coarse-grained density, for instance by being also attracted toward the center of mass of the group. In the future, we plan to study such models coupling the notion of most influential neighbors to a coarse-grained attractive interaction, corresponding to a filtering of the information at the local and global scales, respectively.

In summary, this work provides a comprehensive understanding of the collective dynamics of fish schools in which individuals perform burst-and-coast swimming. We have shown that in this simplified asynchronous burst-and-coast fish school model, both the strength of social interactions and interaction strategies between fish determine the type of collective state. In particular, the strategy which consists for each fish to only interact with its most influential neighbors (*i*.*e*., when *k* = 1), allows the school to display coordinated states over a much larger range of attraction and alignment parameters than when *k* = 2, while still promoting stability of the schooling state over the long term. Finally, the analysis of the long-term dynamics of the schooling, milling, and swarming states, has revealed the typical mode of dispersion that reduces the stability of these states.

## V METHODS

### A Quantification of collective behavior

We characterize the collective behavioral patterns by means of three observables: 1) the group cohesion, measured by the dispersion of individuals with respect to the barycenter of the group, 2) the group polarization, measuring how much fish are aligned, and 3) the milling index, measuring how much the fish turn around their barycenter as in a vortex formation [13].

The *x*-coordinates of the position and velocity vectors of the barycenter *B* are,

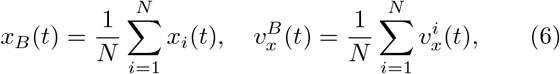

with similar expressions for *y*_*B*_(*t*) and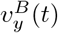. The heading angle of the barycenter is given by its velocity vector,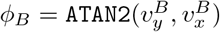. Then:

1. *The group dispersion* D(t) is the mean of the square distance of each fish to the barycenter of the group B:

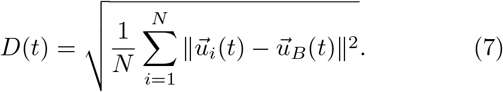 Low values of *D*(*t*) correspond to highly cohesive groups, while high values of *D*(*t*) mean that individuals are spatially dispersed. Values of *D*(*t*) can be arbitrarily high when fish are in a dispersion regime because the distance between fish grows with simulation time.
2. *The group polarization P*(*t*) ∈ [0, 1] is determined by fish headings *ϕ*_*i*_ and independently of the intensity of the speed:

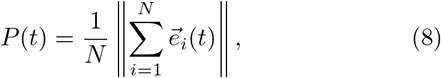

where 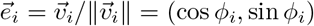 is the unit vector of fish heading. Values of *P* close to 1 mean that the *N* individual headings are aligned and point in the same direction; this is what happens when fish swim in a school formation. Values of *P* close to 0 mean that the *N* vectors point in different directions, but can also mean that vectors are collinear and with opposite direction so that they cancel each other (*e*.*g*., half of the vectors pointing North and the other half pointing South). When headings are uncorrelated, the polarization index is such that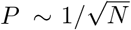, which becomes small only for large group size *N*, but which is markedly lower than 1 for any *N* ≥ 5. For *N* = 100, a value of *P* ≈ 0.1 would mean that the group is not polarized.
3. *The milling index M*(*t*) ∈ [0, 1] measures how much the fish turn in the same direction around their barycenter *B*,

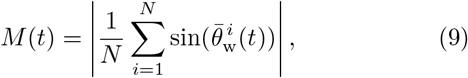

where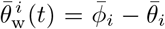. Variables with a bar are defined in the barycenter’s system of reference: 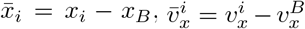 (similar expressions for the y-components).

Then, the relative position angle and heading of fish *i* with respect to *B* are respectively 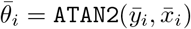 and 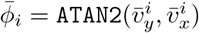. Note that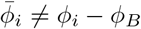.

Fish rotating counterclockwise (resp. clockwise) around *B* make the sum between bars in Eq. (9) to tend to 1 (resp. − 1). The milling index *M*(*t*) denotes how intense is the group milling, whatever the direction of rotation.

### B Numerical simulations of the model

The discretization setting to explore the parameter space was chosen to get a detailed visualization of the regions of interest and their evolution from one condition to another (large phase diagrams were obtained with Δ*γ*_Att_ = 0.005 and Δ*γ*_Ali_ = 0.01.

Parameter values used in the simulations are shown in Table 1*a*. For the color maps of figure 3, we averaged 20 runs of 2000 kicks per fish, and neglected the first half of the time series to remove the effects of the initial condition. For the phase planes of figure 4, we used 20 runs of a longer duration, 2 × 10^4^ kicks per fish, again neglecting the first half. The time series shown in figures 4*e–h* are extracted from the last instants of runs of 2 × 10^4^ kicks per fish. The PDF of figure 5 are the average of 100 runs, each with 2 × 10^4^ kicks. In the long term analysis, we used the average of 100 runs of 2 × 10^4^ kicks per fish in the time series of figure 6, and went to average up to 1000 runs of 4 × 10^4^ kicks per fish for those shown in figure 7.

For every run, the initial conditions were sampled from random distributions and were therefore different. Except for the very long simulations of figures 6 and 7, fish are initially randomly distributed in a circle of radius R chosen so that the mean distance between fish, estimated to be about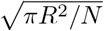, is of the order of half the range of social interactions, so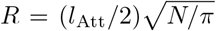. Initial headings are sampled from the uniform distribution in (− *π, π*). For the long term simulations, fish initially have the same heading (*ϕ* = 0) and are distributed in an ellipse of long and short half axes *a* and *b* respectively (long axis taken parallel to the *x*-axis). Values of *a* and *b* for each state and for *k* = 1 and *k* = 2 are shown in Table 1*b*.

### C Calculation of the number of groups

The number of groups *N*_G_ is calculated with the “chain of offspring” recursive method. We first consider that two fish that are separated by more than a critical distance of interaction do not interact with each other. We tried several critical distances and found that using a distance of four times the interaction range of attraction the re-sult is in agreement with the direct observation of the spatial configuration. Consequently, an individual fish whose nearest neighbor is further than this critical distance constitutes a group (of size 1) in itself. The rest of groups are then build as follows. Starting with a given fish, add to the group of this fish its nearest neighbor, the nearest neighbor of its nearest neighbor, and so on, until the next nearest neighbor is already in the group. Then, another fish is taken that is not yet in a group, and its “family of offspring” is built with the same recursive procedure.

It may happen that one or more groups are in fact at a close distance one from each other, or even that a group is located in the interior of another group. In that case, individuals from one group can be subject to the influence of individuals of another group, so that the fish are not really in different groups. To prevent these situations, we merge the groups that are separated by a distance shorter than the minimum interaction range, min{*l*_Att_, *l*_Ali_}.

#### Ethics

This work did not require ethical approval from a human subject or animal welfare committee.

## Authors contribution

G.T. designed research; W.W. and R.E. performed research; W.W., R.E., C.S. and G.T. analyzed data; W.W. performed numerical simulations. W.W., R.E. and G.T. wrote the paper with input from all other authors.

## Acknowledgements

G.T. acknowledge the support of the Indo-French Centre for the Promotion of Advanced Research (Project No. 64T4-B). G.T. also gratefully acknowledges the Indian Institute of Science for support via Infosys Visiting Chair Professor at the Centre for Ecological Sciences, IISc, Bengaluru.

